# The association between 25(OH)D levels, frailty status and adiposity indices in older adults

**DOI:** 10.1101/330340

**Authors:** Ana Rita Sousa-Santos, Cláudia Afonso, Alejandro Santos, Nuno Borges, Pedro Moreira, Patrícia Padrão, Isabel Fonseca, Teresa F Amaral

## Abstract

**Background:** Vitamin D deficiency is common in older adults and has been linked with frailty and obesity, but it remains to be studied whether frail obese older adults are at higher risk of vitamin D deficiency. Therefore, the aim of this study is to explore the association between frailty, adiposity indices and serum 25(OH)D concentrations.

**Methods:** 1447 individuals with 65 years or older, participating in a cross-sectional study (Nutrition UP 65). Frailty, according to Fried et al., body mass index (BMI), waist circumference (WC), body roundness index (BRI) and body shape index (ABSI) were evaluated. A stepwise multinomial logistic regression was carried out to quantify the association between 25(OH)D quartiles and independent variables.

**Results:** Median 25(OH)D levels were lower in individuals presenting both frailty and obesity (*p*<0.001). In the multivariate analysis, pre-frailty (OR 2.65; 95% CI 1.63-4.32) and frailty (OR 3.76; 95% CI 2.08-6.81) were associated with increased odds of lower 25(OH)D serum levels (first quartile). Regarding adiposity indices, obesity (OR 1.75; 95% CI 1.07-2.87) and the highest categories of WC (OR 3.46; 95% CI 1.95-6.15), BRI (OR 4.35; 95% CI 2.60-7.29) and ABSI (OR 3.17 95% CI 1.86-5.38) were directly associated with lower 25(OH)D serum levels (first quartile).

**Conclusions:** A positive association between frailty or obesity and lower levels of vitamin D was found. Moreover, besides BMI and WC, other indicators of body adiposity, such as BRI and ABSI, were associated with lower 25(OH)D serum concentrations.

## Introduction

Vitamin D is fat-soluble vitamin mainly obtained from sun exposure of the skin and in lesser amounts from diet and supplements (1–3). It is stored mainly in adipose tissue and muscle and, to a lesser extent, in other tissues (4). Vitamin D deficiency is a public health problem of growing concern (5–7), common in older adults (5,7,8) and it has been linked to adverse health outcomes such as falls (9), poorer cognitive function (10) and cancer (11). 25(OH)D concentrations decrease with age, due to a reduction in cutaneous vitamin D synthesis (12), to the possible decline in the ability of the kidney to synthesize 1,25(OH)_2_D (4).

Despite the well-known consequences of vitamin D deficiency in bone health (13), this hormone seems to also have a key-role in skeletal muscle (14), namely influencing its function and performance (14,15). Frailty increases with age and its prevalence in the community ranges from 4.0-59.1%, depending on the definition adopted (16). It is associated with an increased risk of adverse health outcomes, such as falls, disability, hospitalization and even mortality (17). Evidence has shown a link between frailty and vitamin D status, with frailty being associated with lower levels of serum 25(OH)D (18). However, the impact of vitamin D deficiency in frailty status in later life is still unknown.

Obesity has also increased appreciably worldwide and older adults are no exception (19). Several meta-analyses reported a significant association with lower serum 25(OH)D concentrations (20–22), although the mechanisms underlying this association are not yet fully understood. Furthermore, obesity has also been positively associated with frailty status in older adults (23,24), but it remains to be studied whether frail obese older adults are at higher risk of vitamin D deficiency and if the presence of these conditions could simultaneously lead to worse health outcomes. According to the previously described in literature, obese older adults may be predisposed to vitamin D deficiency, which is in turn associated with worse physical function and frailty (18,25). Conversely, frailty may impact the amount of sun exposure and, consequently, predispose to vitamin D deficiency. Even though several studies have evaluated the association of frailty status and obesity on vitamin D levels separately (18,20,21), to our knowledge, literature regarding the study of all three conditions is absent. It will be relevant to know if frail obese older adults are more likely to present low vitamin D levels. Besides body mass index (BMI), other adiposity indicators such as waist circumference (WC), body roundness index (BRI) and body shape index (ABSI) may be used (26,27). While previous studies have established a link between several indices and vitamin D status, such as BMI and WC (28,29), data regarding BRI and ABSI is lacking. Thus, we hypothesized that these indices may be associated with vitamin D status, as higher values may denote worse vitamin D status.

Frailty, obesity and vitamin D deficiency are potentially preventable or treatable. Early interventions in these conditions may lead to an improvement in health status and quality of life during the course of ageing (30). So, it is important to elucidate the association between these conditions to target the individuals at risk. Therefore, the aim of this study is to evaluate the association between serum 25(OH)D concentrations, frailty and obesity, but also to examine if there is an interaction effect between frailty and obesity on 25(OH)D levels. In addition, the association of other adiposity indicators, such as WC, BRI and ABSI, with vitamin D status was also explored.

## Materials and Methods

The study sample included individuals enrolled in the Nutrition UP 65 study, a cross-sectional observational study conducted in Portugal. As described in detail previously (31), a cluster sample of 1500 individuals with 65 years or older, representative of the Portuguese older population in terms of age, sex, education and regional area was selected. In each regional area, three or more town councils with >250 inhabitants were randomly selected and potential community-dwelling participants were contacted via home approach, telephone or via institutions such as town councils and parish centres. Individuals presenting any condition that precluded the collection of venous blood samples or urine (*eg,* dementia or urinary incontinence) were not included.

Data were gathered between December 2015 and June 2016. A structured questionnaire was applied by interview, conducted by eight trained registered nutritionists and anthropometric data were also collected. From the initial sample, forty-six individuals could not be assessed regarding frailty status (n=43) and body mass index (n=4) due to missing data and were therefore excluded from the present analysis. Additionally, seven older adults were also excluded due to missing data regarding the covariates.

## Anthropometric and functional measurements

Anthropometric measurements were collected following standard procedures (32). A calibrated stadiometer (SECA 213, SECA GmbH, Hamburg, Germany) with 0.1 cm resolution was used to measure standing height. Body weight (in kilograms) was measured with a calibrated portable electronic scale (SECA 803, SECA GmbH, Hamburg, Germany) with 0.1 kg resolution, with the participants wearing light clothes. When it was not possible to measure standing height or weigh a participant height was obtained indirectly from non-dominant hand length (32), measured with a calibrated caliper (Fervi Equipment) with 0.1 centimeter resolution and body weight was estimated from mid-upper arm and calf circumferences (33). Mid upper arm, waist and calf circumferences were measured with a metal tape measure (Lufkin W606 PM, Lufkin^®^, Sparks, Maryland, USA) with 0.1 cm resolution. Triceps skinfold thickness was obtained using a Holtain Tanner/Whitehouse (Holtain, Ltd., Crosswell, United Kingdom) skinfold calliper with 0.2 mm resolution.

Hand grip strength (HGS) was measured in the non-dominant hand with a calibrated Jamar Plus Digital Hand Dynamometer (Sammons Preston Inc., Bolingbrook, Illinois, USA). As recommended by the American Society of Hand Therapists, participants were asked to sit in a chair without arm rest, with their shoulders adducted, their elbows flexed 90° and their forearms in neutral position (34). Three measurements with a one-minute pause between them were performed by each individual and the higher value, recorded in kilogram-force (kgf), was used for the analysis. Individuals unable to perform the measurement with the non-dominant hand were asked to use the dominant hand.

Walking time was measured over a distance of 4.6 meters, in an unobstructed corridor. Individuals were instructed to walk at usual pace and walking time was recorded by a chronometer (School electronic stopwatch, Dive049, Topgim, Portugal), in seconds. Those unable to perform the test due to mobility or balance limitations were considered frail for this criterion (n=28).

Self-reported exhaustion was measured using two items from the Center for Epidemiologic Studies Depression Scale (CES-D) (35). The following two statements were read: “I felt that everything I did was an effort” and “In the last week I could not get going.” The exhaustion criterion was considered present if a participant answered “a moderate amount of the time” or “most of the time” to the question: “How often in the last week did you feel this way?”.

Physical activity, assessed by the short form of the International Physical Activity Questionnaire (36), included information regarding the previous seven days, namely on how many days and how much time the participant spent: walking or hiking (at home or at work, moving from place to place, for recreation or sport), sitting (at a desk, visiting friends, reading, studying or watching television), moderate activities (carrying light objects, hunting, carpentry, gardening, cycling at a normal pace or tennis in pairs) and vigorous activities, namely lifting heavy objects, agriculture, digging, aerobics, swimming, playing football and cycling at a fast pace was gathered.

## Frailty status

Frailty was defined according to Fried *et al.* frailty phenotype (17). Pre-frailty was classified as the presence of one or two of criteria and frailty as the presence of three or more of the following five criteria: “shrinking”: evaluated by self-reported unintentional weight loss (>4.5 kg lost unintentionally in prior year); “weakness”: assessed by low HGS adjusted for sex and BMI; “poor endurance and energy”: evaluated by self-reported exhaustion; “slowness”: identified by walking time adjusted for sex and standing height and “low physical activity”: by means of energy expended per week, adjusted for sex (men <383 kcal/week and women <270 kcal/week).

## Laboratory analyses

Qualified nurses collected blood samples for these analyses, preferentially after a 12-hour fasting period. Vitamin D status was evaluated by dosing the plasmatic levels of 25-hydroxycholecalciferol through the electrochemiluminescence immunoassay using Roche Cobas Vitamin D total assay reagent (Roche Diagnostics GmbH, Mannheim, Germany). All samples were analysed with the same equipment. Since 25(OH)D serum concentrations were very low in our sample, 25(OH)D concentrations were categorized into quartiles (Q). For characterization purposes individuals were still classified according to the Institute of Medicine (IOM) criteria as being at risk of deficiency at serum 25(OH)D concentrations <12 ng/mL, at risk for inadequacy at levels ranging from 12–<20 ng/mL and having sufficient at levels when 25(OH)D concentrations are ≥20 ng/mL (37). Data concerning 25(OH)D levels in Nutrition UP 65 study was previously described (8,38).

## Adiposity indices

BMI was calculated as (weight (kg)/ height^2^ (m)), and subjects were classified as underweight for BMI below 18.5 kg/m^2^, as normal weight for BMI between 18.5-24.9 kg/m^2^, as pre-obese for BMI between 25.0-29.9 kg/m^2^ and as obese for BMI above 30.0 kg/m^2^ (39). Underweight individuals were included in the reference group (“normal weight”) due to its small number (n=3). WC was categorized according to the risk of metabolic complications as increased (men >94 cm; women >80 cm) and substantially increased (men >102 cm; women >88 cm) (40). BRI was calculated based on WC (m) and height (m) (27):

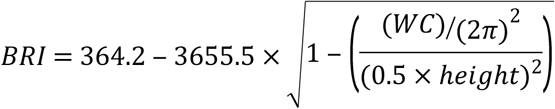

ABSI (m^11/6^·kg^−2/3^) was calculated according to the following formula, based on WC (m), BMI (kg/m^2^) and height (m) (26):

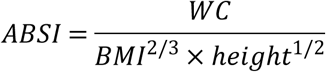

Quartiles of BRI and ABSI were calculated.

## Variables collection and categorization

Information regarding educational level, smoking status, alcohol consumption and vitamin D supplementation was self-reported. Educational level was determined by the number of completed school years and the following categories were used: without schooling, 1-4 years, 5-12 years and >12 years. All individuals reported information on smoking status and this information was included as a dichotomous variable: smoker or non-smoker. Alcohol consumption was evaluated as the number of alcoholic drinks daily and was included in the analyses as a categorical variable: none, moderate consumption (women ≤ 1 and men ≤ 2 alcoholic drink daily), and excessive consumption (women > 1 and men > 2 alcoholic drinks daily). The Portuguese version of the Mini Mental State Examination was used to ascertain cognitive decline, which was dichotomized into not impaired and impaired. Cut-off scores for cognitive impairment were the following: individuals with no education, ≤15 points; 1 to 11 years of years of school completed, ≤22 points; and >11 years of school completed, ≤27 points (41). Season of blood collection was presented as a dichotomous variable: spring/summer or autumn/winter. Skin phenotype was defined according to the Fitzpatrick classification: red-haired with freckles, fair-haired, dark-haired, Latin, Arab, Asian or Black (42). Vitamin D supplements use was categorized as: no use, use of vitamin D supplements, unknown composition or use.

## Ethics

This research was conducted according to the guidelines established by the Declaration of Helsinki, and the study protocol was approved by the Ethics Committee of the Department of “Ciências Sociais e Saúde” (Social Sciences and Health) from the “Faculdade de Medicina da Universidade do Porto” (PCEDCSS – FMUP 15/2015) and by the Portuguese National Commission of Data Protection (9427/2015). All study participants signed an informed consent form.

## Statistical analyses

Descriptive analyses were conducted to compare participants’ characteristics across 25(OH)D quartiles. Results were presented as number of participants (expressed as percentage), for categorical variables. For continuous variables, means (standard deviations) were used, or medians (interquartile range) to report variables with skewed distribution. ANOVA or Kruskal-Wallis were used to test associations between the study groups and continuous variables. Multiple comparisons between frailty and obesity groups were performed using Dunn-Bonferroni tests. Differences in proportions, as well comparison between included and excluded individuals, in the sensitivity analysis, were tested using Chi-square test or Fisher’s exact test.

A multinomial logistic regression was carried out to quantify the association between 25(OH)D quartiles (dependent variable) and independent variables. Odds ratios (OR) and their respective 95% confidence intervals (CI) were calculated, with adjustments for sex, age, smoking status, alcohol consumption, season of blood collection, vitamin D supplementation and skin phenotype. A stepwise approach with forward entry was carried out to explore the following interactions terms in each model: frailty status*BMI, frailty status*WC, frailty status*BRI and frailty status*ABSI.

Statistical significance was established at a *p*-value <0.05. All statistical analyses were conducted with IBM SPSS Statistics 23 (SPSS, Inc, an IBM Company, Chicago, IL).

## Results

Descriptive data of the 1447 older adults (57.8% women) included in this study and statistical differences in sociodemographic lifestyle and health conditions according to 25(OH)D quartiles are shown in Table 1. Median age of the individuals was 74 years (range 65-100). Based on Fried’s frailty definition, 21.4% were frail and 39% were obese according to BMI. Overall, the majority of older adults were non-smokers, however slightly more than half (51.3%) reported consuming alcoholic drinks daily.

**Table 1.**
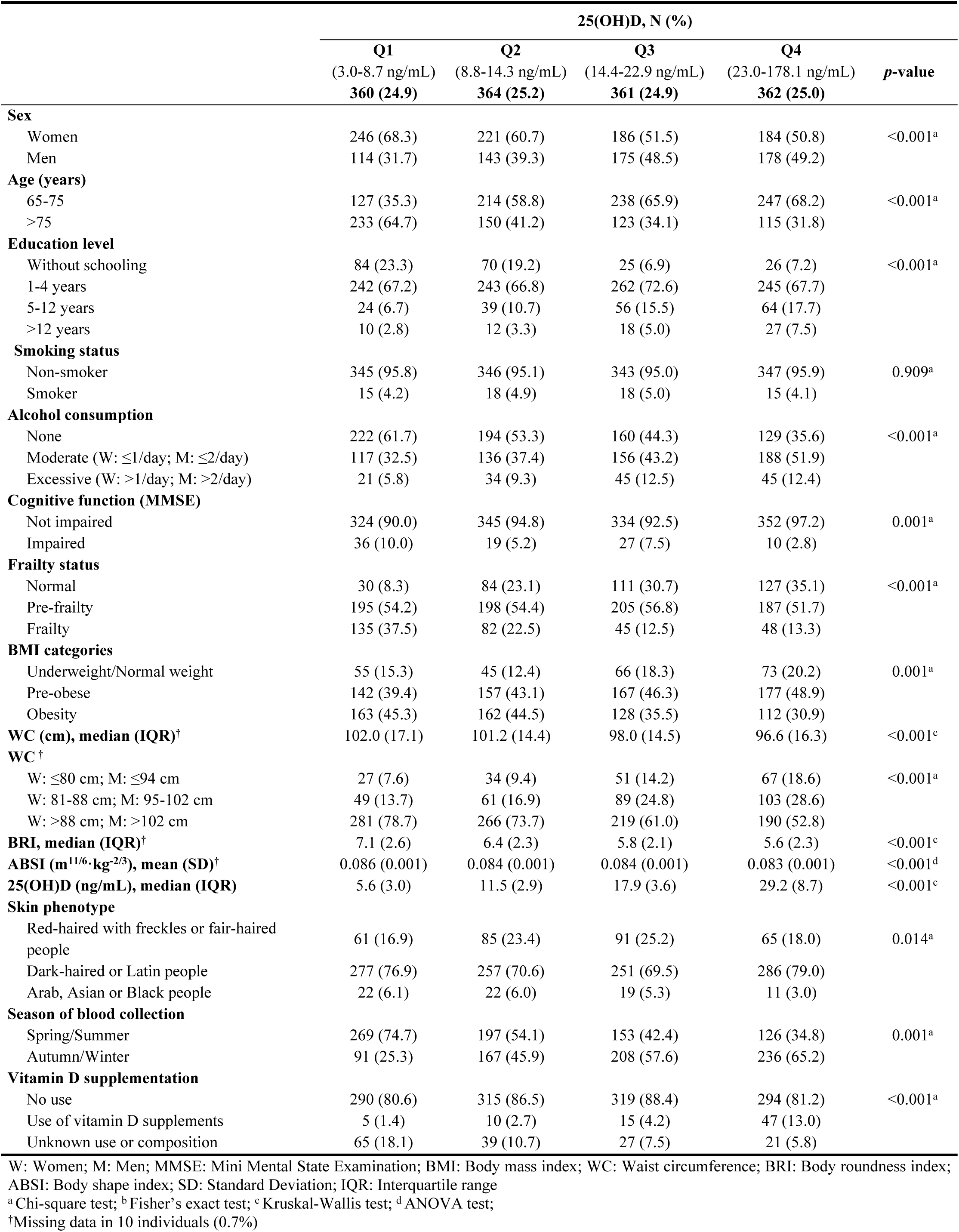
Characteristics of the 1447 included older adults by 25(OH)D quartiles.

Regarding vitamin D serum levels, 69% of participants had 25(OH)D <20 ng/mL, and 39.7% had 25(OH)D <12 ng/mL. Additionally, only 5.3% reported the use of vitamin D supplements. Median 25(OH)D levels of Q1 were 5.6 ng/mL (interquartile range (IQR): 3.0 ng/mL), for Q2 were 11.5 ng/mL (IQR: 2.9 ng/mL), for Q3 were 17.9 ng/mL (IQR: 3.6 ng/mL) and, lastly, for Q4 were 29.2 (IQR: 8.7 ng/mL). When studying participants’ characteristics according to 25(OH)D quartiles, significant differences were observed for all studied variables, except for smoking status. As expected, individuals that reported the use vitamin D supplements were more likely to present higher 25(OH)D serum values and to fit in the fourth quartile (*p*<0.001). Moreover, a higher proportion of frail (*p*<0.001), obese (*p*=0.001) and cognitive impaired (*p*=0.001) older adults was observed in the first quartile of 25(OH)D levels. Median and mean values of WC, BRI and ABSI decreased across increased quartiles of 25(OH)D levels (*p*<0.001).

Sensitivity analysis comparing excluded and included older adults in the present study, showed that those who were excluded reported a lower alcohol consumption (*p*=0.001), were more likely to be cognitively impaired (*p*=0.049) and to have a darker skin phenotype (*p*=0.002) (Supplementary Table 1).

Regarding coexistence of frailty and obesity (Supplementary Table 2), approximately 75% of older adults presenting both frailty and obesity were women and nearly 67% were aged over 75 years. More than 60% of participants with one at least condition (either frailty or obesity or both) were women (*p*≤0.001). Individuals presenting only obesity were more likely to be younger (64.9%) and 70.9% of the older adults presenting only frailty were in the oldest category (*p*<0.001). Median 25(OH)D values decreased across obesity and frailty status and were 17.1 ng/mL for non-obese non-frail individuals, 13.6 ng/mL for obese non-frail individuals, 10.1 ng/mL for non-obese frail individuals and 9.2 ng/mL in individuals presenting both obesity and frailty (*p*<0.001) (Supplementary Table 2).

To evaluate the association between the indices BMI, WC, BRI and ABSI, Spearman correlation coefficients were also calculated. BMI was positively and significantly correlated with WC (ρ=0.748), BRI (ρ=0.842) and negatively correlated with ABSI (ρ=-0.121). However, ABSI correlated positively and significantly with WC (ρ=0.476) and BRI (ρ=0.358) (Supplementary Table 3).

Comparisons of median 25(OH)D serum levels between obesity and frailty status groups, are displayed in Figure 1. Median 25(OH)D levels in all groups were below 20 ng/ml. In women, median 25(OH)D levels were significantly higher in non-obese non-frail group, comparing with obese non-frail group [15.7 (IQR: 16.5) vs 12.4 (IQR: 16.5)], *p*=0.007; non-obese frail group [15.7 (IQR: 16.5) vs 10.2 (IQR: 11.9)], *p*<0.001 and obese frail group [15.7 (IQR: 16.5) vs 9.0 (IQR: 11.0)], p<0.001. Also, obese non-frail women had significantly higher median 25(OH)D levels comparing with the obese frail group [12.4 (IQR: 16.5) vs 9.0 (IQR: 11.0)], *p*=0.006. Among men, median 25(OH)D levels were only significantly higher in non-obese non-frail group, comparing with non-obese frail [17.7 (IQR: 14.1) vs 9.5 (IQR: 9.2)] and obese frail group [17.7 (IQR: 14.1) vs 10.1 (IQR: 10.8)], *p*<0.001. However, the obese non-frail group also presented significantly higher median 25(OH)D levels than non-obese frail [18.2 (IQR: 15.0) vs 9.5 (IQR: 9.2)] and obese frail groups [18.2 (IQR: 15.0) vs 10.1 (IQR: 10.8)], *p*<0.001 and *p*=0.002, respectively.

**Fig 1.**
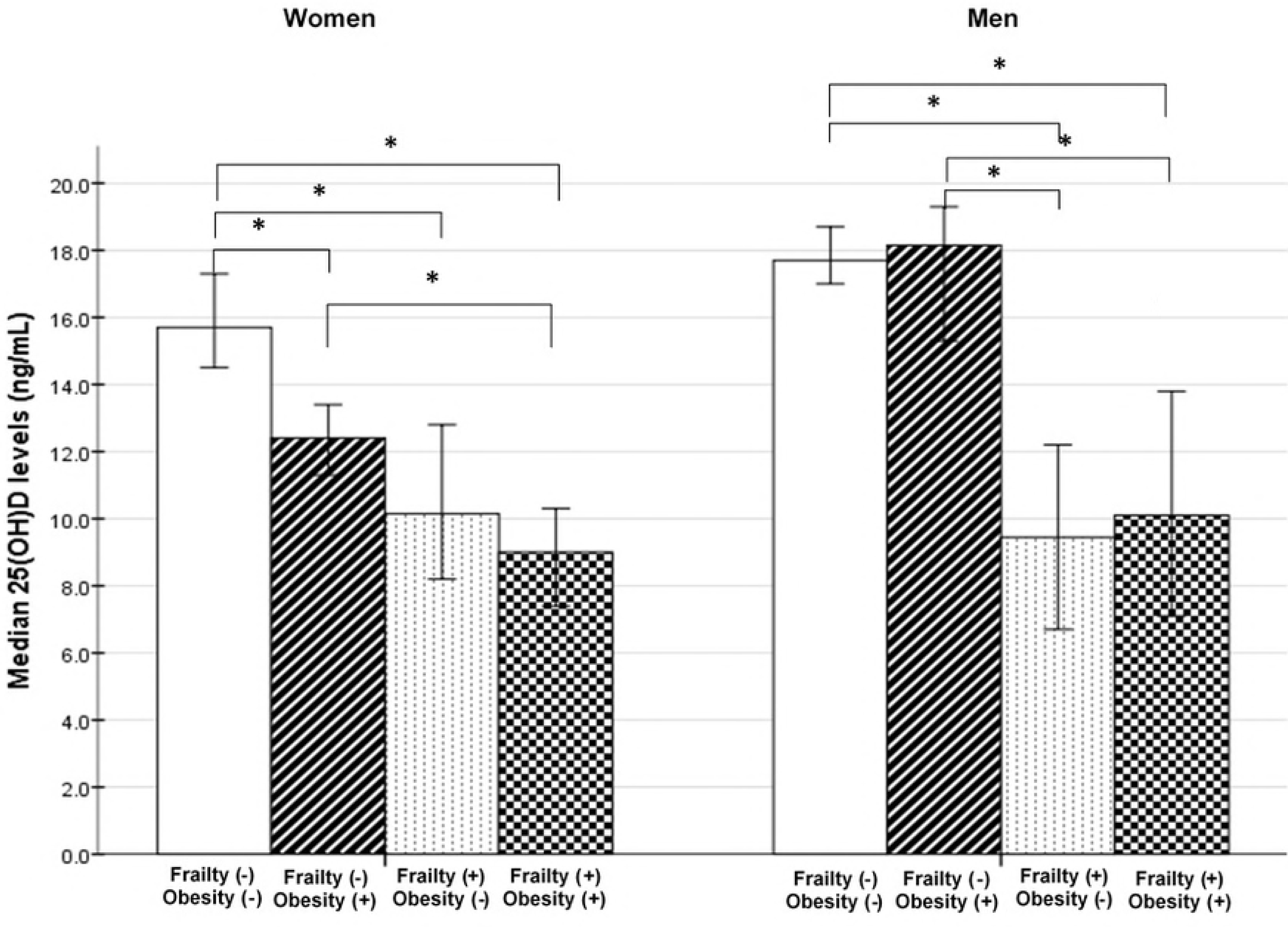
Differences in median (95% CI) 25(OH)D serum levels between Obesity(-) Frailty(-) (W: n=359; M: n=365), Obesity(+) Frailty(-) (W: n=263; M: n=150), Obesity(-) Frailty(+) (W: n=102; M: n=56) and Obesity(+) Frailty(+) (W: n=113; M: n=39) groups, in older women (W) and men (M), using Kruskal-Wallis with Dunn-Bonferroni tests for multiple comparisons. *p<0.05 for pairwise comparisons.

The association between obesity and frailty status and 25(OH)D quartiles was further investigated through multivariate multinomial regression (Table 2). Considering the fourth quartile of serum 25(OH)D as the reference category, pre-frail older adults were 2.65 (95% CI: 1.63-4.32) times more likely to be in the first quartile of serum 25(OH)D (3.0-8.7 ng/mL), and frail individuals were 3.76 (95% CI: 2.08-6.81) times more likely to present serum 25(OH)D levels in the first quartile (*P* for trend <0.001). For individuals in the two lowest quartiles of serum 25(OH)D levels (Q1: 3.0-8.7 ng/mL and Q2: 8.8-14.3 ng/mL), the adjusted odds ratios for obesity were 1.75 (95% CI: 1.07-2.87) for the first (*P* for trend=0.011) and 2.21 (95% CI: 1.37-3.55) second (*P* for trend <0.001) quartiles, respectively. The association between pre-obesity and serum 25(OH)D levels did not reach statistical significance in any quartile.

**Table 2.**
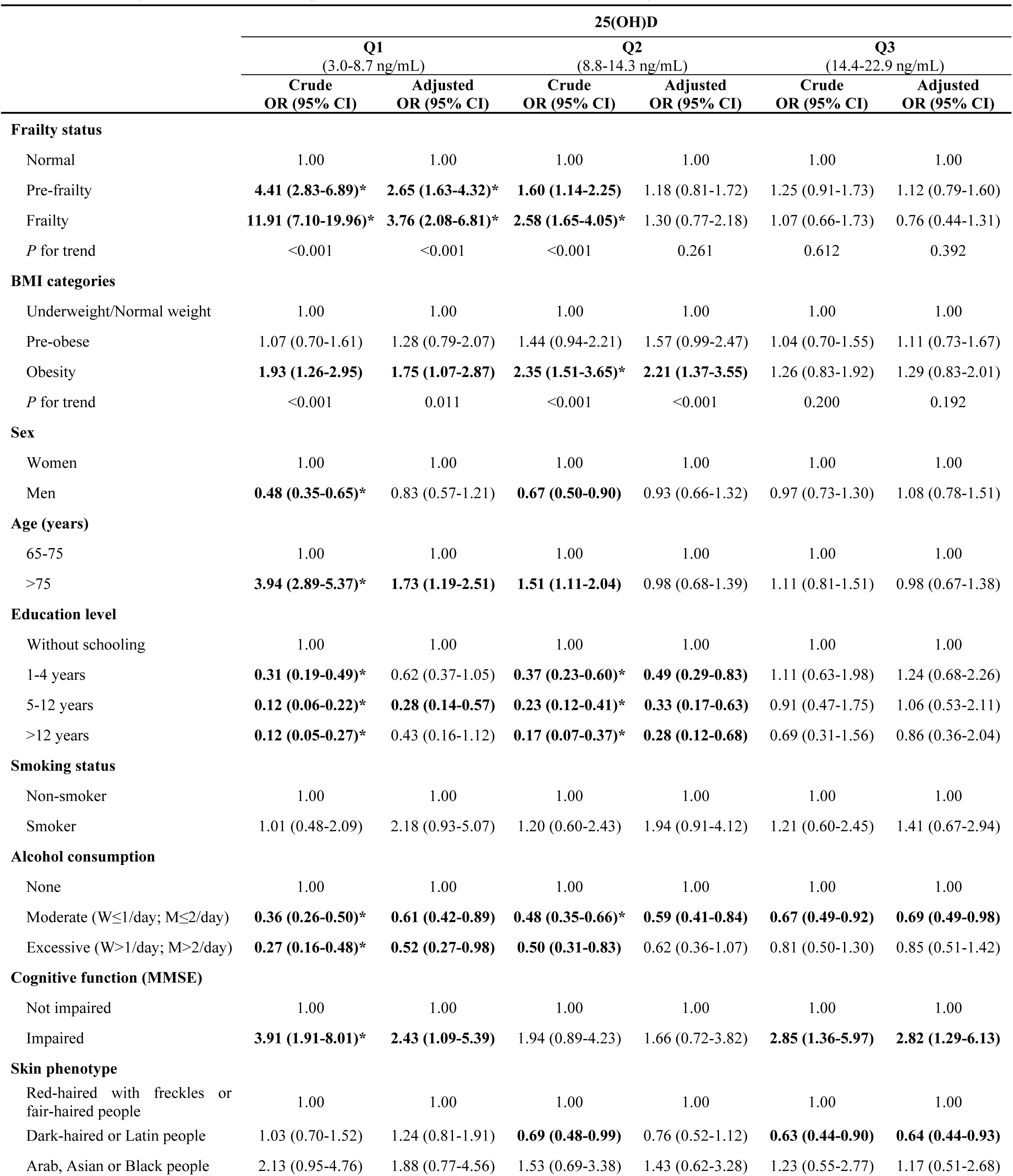
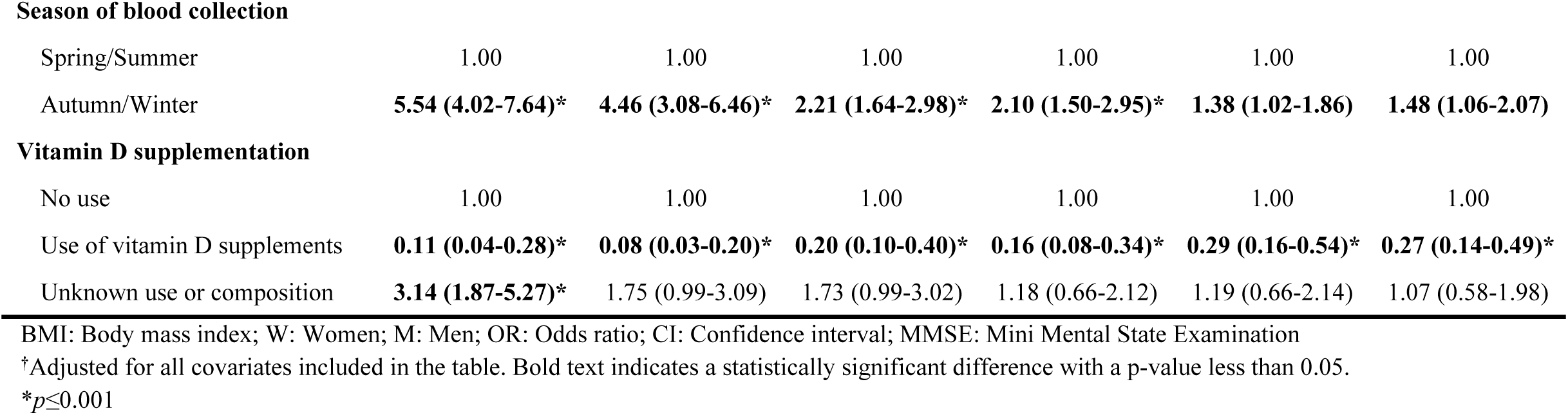
Multinomial logistic regression regarding frailty status and body mass index with 25(OH)D quartiles. Reference category was the fourth quartile of serum 25(OH)D (23.0-178.1 ng/mL)^†^.

|  | 25(OH)D |  |  |  |  |  |
| --- | --- | --- | --- | --- | --- | --- |
|  | Q1 |  | Q2 |  | Q3 |  |
|  | (3.0-8.7 ng/mL) |  | (8.8-14.3 ng/mL) |  | (14.4-22.9 ng/mL) |  |
|  | Crude<br>OR (95% CI) | Adjusted<br>OR (95% CI) | Crude<br>OR (95% CI) | Adjusted<br>OR (95% CI) | Crude<br>OR (95% CI) | Adjusted<br>OR (95% CI) |
| **Frailty status** |  |  |  |  |  |  |
| Normal | 1.00 | 1.00 | 1.00 | 1.00 | 1.00 | 1.00 |
| Pre-frailty | **4.41 (2.83-6.89)\*** | **2.65 (1.63-4.32)\*** | **1.60 (1.14-2.25)** | 1.18 (0.81-1.72) | 1.25 (0.91-1.73) | 1.12 (0.79-1.60) |
| Frailty | **11.91 (7.10-19.96)\*** | **3.76 (2.08-6.81)\*** | **2.58 (1.65-4.05)\*** | 1.30 (0.77-2.18) | 1.07 (0.66-1.73) | 0.76 (0.44-1.31) |
| *P* for trend | <0.001 | <0.001 | <0.001 | 0.261 | 0.612 | 0.392 |
| **BMI categories** |  |  |  |  |  |  |
| Underweight/Normal weight | 1.00 | 1.00 | 1.00 | 1.00 | 1.00 | 1.00 |
| Pre-obese | 1.07 (0.70-1.61) | 1.28 (0.79-2.07) | 1.44 (0.94-2.21) | 1.57 (0.99-2.47) | 1.04 (0.70-1.55) | 1.11 (0.73-1.67) |
| Obesity | **1.93 (1.26-2.95)** | **1.75 (1.07-2.87)** | **2.35 (1.51-3.65)\*** | **2.21 (1.37-3.55)** | 1.26 (0.83-1.92) | 1.29 (0.83-2.01) |
| *P* for trend | <0.001 | 0.011 | <0.001 | <0.001 | 0.200 | 0.192 |
| **Sex** |  |  |  |  |  |  |
| Women | 1.00 | 1.00 | 1.00 | 1.00 | 1.00 | 1.00 |
| Men | **0.48 (0.35-0.65)\*** | 0.83 (0.57-1.21) | **0.67 (0.50-0.90)** | 0.93 (0.66-1.32) | 0.97 (0.73-1.30) | 1.08 (0.78-1.51) |
| **Age (years)** |  |  |  |  |  |  |
| 65-75 | 1.00 | 1.00 | 1.00 | 1.00 | 1.00 | 1.00 |
| >75 | **3.94 (2.89-5.37)\*** | **1.73 (1.19-2.51)** | **1.51 (1.11-2.04)** | 0.98 (0.68-1.39) | 1.11 (0.81-1.51) | 0.98 (0.67-1.38) |
| **Education level** |  |  |  |  |  |  |
| Without schooling | 1.00 | 1.00 | 1.00 | 1.00 | 1.00 | 1.00 |
| 1-4 years | **0.31 (0.19-0.49)\*** | 0.62 (0.37-1.05) | **0.37 (0.23-0.60)\*** | **0.49 (0.29-0.83)** | 1.11 (0.63-1.98) | 1.24 (0.68-2.26) |
| 5-12 years | **0.12 (0.06-0.22)\*** | **0.28 (0.14-0.57)** | **0.23 (0.12-0.41)\*** | **0.33 (0.17-0.63)** | 0.91 (0.47-1.75) | 1.06 (0.53-2.11) |
| >12 years | **0.12 (0.05-0.27)\*** | 0.43 (0.16-1.12) | **0.17 (0.07-0.37)\*** | **0.28 (0.12-0.68)** | 0.69 (0.31-1.56) | 0.86 (0.36-2.04) |
| **Smoking status** |  |  |  |  |  |  |
| Non-smoker | 1.00 | 1.00 | 1.00 | 1.00 | 1.00 | 1.00 |
| Smoker | 1.01 (0.48-2.09) | 2.18 (0.93-5.07) | 1.20 (0.60-2.43) | 1.94 (0.91-4.12) | 1.21 (0.60-2.45) | 1.41 (0.67-2.94) |
| **Alcohol consumption** |  |  |  |  |  |  |
| None | 1.00 | 1.00 | 1.00 | 1.00 | 1.00 | 1.00 |
| Moderate (W≤1/day; M≤2/day) | **0.36 (0.26-0.50)\*** | **0.61 (0.42-0.89)** | **0.48 (0.35-0.66)\*** | **0.59 (0.41-0.84)** | **0.67 (0.49-0.92)** | **0.69 (0.49-0.98)** |
| Excessive (W>1/day; M>2/day) | **0.27 (0.16-0.48)\*** | **0.52 (0.27-0.98)** | **0.50 (0.31-0.83)** | 0.62 (0.36-1.07) | 0.81 (0.50-1.30) | 0.85 (0.51-1.42) |
| **Cognitive function (MMSE)** |  |  |  |  |  |  |
| Not impaired | 1.00 | 1.00 | 1.00 | 1.00 | 1.00 | 1.00 |
| Impaired | **3.91 (1.91-8.01)\*** | **2.43 (1.09-5.39)** | 1.94 (0.89-4.23) | 1.66 (0.72-3.82) | **2.85 (1.36-5.97)** | **2.82 (1.29-6.13)** |
| **Skin phenotype** |  |  |  |  |  |  |
| Red-haired with freckles or fair-haired people | 1.00 | 1.00 | 1.00 | 1.00 | 1.00 | 1.00 |
| Dark-haired or Latin people | 1.03 (0.70-1.52) | 1.24 (0.81-1.91) | **0.69 (0.48-0.99)** | 0.76 (0.52-1.12) | **0.63 (0.44-0.90)** | **0.64 (0.44-0.93)** |
| Arab, Asian or Black people | 2.13 (0.95-4.76) | 1.88 (0.77-4.56) | 1.53 (0.69-3.38) | 1.43 (0.62-3.28) | 1.23 (0.55-2.77) | 1.17 (0.51-2.68) |
| Season of blood collection |  |  |  |  |  |  |
| Spring/Summer | 1.00 | 1.00 | 1.00 | 1.00 | 1.00 | 1.00 |
| Autumn/Winter | **5.54 (4.02-7.64)\*** | **4.46 (3.08-6.46)\*** | **2.21 (1.64-2.98)\*** | **2.10 (1.50-2.95)\*** | **1.38 (1.02-1.86)** | **1.48 (1.06-2.07)** |
| **Vitamin D supplementation** |  |  |  |  |  |  |
| No use | 1.00 | 1.00 | 1.00 | 1.00 | 1.00 | 1.00 |
| Use of vitamin D supplements | **0.11 (0.04-0.28)\*** | **0.08 (0.03-0.20)\*** | **0.20 (0.10-0.40)\*** | **0.16 (0.08-0.34)\*** | **0.29 (0.16-0.54)\*** | **0.27 (0.14-0.49)\*** |
| Unknown use or composition | **3.14 (1.87-5.27)\*** | 1.75 (0.99-3.09) | 1.73 (0.99-3.02) | 1.18 (0.66-2.12) | 1.19 (0.66-2.14) | 1.07 (0.58-1.98) |

BMI: Body mass index; W: Women; M: Men; OR: Odds ratio; CI: Confidence interval; MMSE: Mini Mental State Examination
†Adjusted for all covariates included in the table. Bold text indicates a statistically significant difference with a p-value less than 0.05.
\* $p \leq 0.001$

Multinomial logistic regressions were conducted to evaluate the association of WC, BRI and ABSI with serum 25(OH)D using 25(OH)D quartiles as the dependent variable and the fourth quartile as the reference category (Table 3). Older adults in the highest category of WC presented the odds of 3.46 (95% CI 1.95-6.15) and 2.61 (95% CI 1.58-4.29) for being in the first and second serum 25(OH)D levels quartiles, respectively (*P* for trend <0.001). Although no specific significant associations were identified in the third 25(OH)D quartile, a significant trend was also observed (*P* for trend=0.041). The participants in first quartile of serum 25(OH)D levels showed an increasing adjusted odds ratio for BRI, from the second through the fourth quartile: 1.69 (95% CI: 1.02-2.79), 2.26 (95% CI: 1.36-3.75) and 4.35 (95% CI: 2.60-7.29), (*P* for trend <0.001). Regarding ABSI and for the participants placed in the lowest 25(OH)D serum levels (first) quartile, the odds ratios were 4.03 (95% CI: 2.37-7.86) and 3.17 (95% CI: 1.86-5.38) for the third and fourth ABSI quartiles, respectively (*P* for trend <0.001).

**Table 3.** Association between waist circumference, body roundness index and body shape index with 25(OH)D quartiles. Multinomial logistic regression models. Reference category was the fourth quartile of serum 25(OH)D (23.0-178.1 ng/mL)^†^.

|  | 25(OH)D |  |  |  |  |  |
| --- | --- | --- | --- | --- | --- | --- |
|  | Q1<br>(3.0-8.7 ng/mL) |  | Q2<br>(8.8-14.3 ng/mL) |  | Q3<br>(14.4-22.9 ng/mL) |  |
|  | Crude<br>OR (95% CI) | Adjusted<sup>†</sup><br>OR (95% CI) | Crude<br>OR (95% CI) | Adjusted<sup>†</sup><br>OR (95% CI) | Crude<br>OR (95% CI) | Adjusted<sup>†</sup><br>OR (95% CI) |
| **Waist circumference (WC)**□ |  |  |  |  |  |  |
| W: ≤80cm; M: ≤94cm | 1.00 | 1.00 | 1.00 | 1.00 | 1.00 | 1.00 |
| W: 81-88 cm; M: 95-102 cm | 1.18 (0.67-2.07) | 1.31 (0.69-2.48) | 1.17 (0.69-1.96) | 1.27 (0.73-2.20) | 1.14 (0.72-1.80) | 1.23 (0.76-1.98) |
| W: >88 cm; M: >102 cm | **3.67 (2.26-5.95)\*** | **3.46 (1.95-6.15)\*** | **2.76 (1.75-4.34)\*** | **2.61 (1.58-4.29)\*** | **1.51 (1.00-2.29)** | 1.56 (0.99-2.45) |
| P for trend | <0.001 | <0.001 | <0.001 | <0.001 | 0.019 | 0.041 |
| **Body roundness index (BRI)**□ |  |  |  |  |  |  |
| Q1 (1.93-5.09) | 1.00 | 1.00 | 1.00 | 1.00 | 1.00 | 1.00 |
| Q2 (5.10-6.10) | **1.78 (1.14-2.78)** | **1.69 (1.02-2.79)** | **1.68 (1.12-2.52)** | 1.51 (0.98-2.31) | 1.27 (0.87-1.86) | 1.15 (0.77-1.71) |
| Q3 (6.11-7.49) | **2.88 (1.85-4.50)\*** | **2.26 (1.36-3.75)** | **2.48 (1.64-3.75)\*** | **2.14 (1.37-3.34)** | **1.56 (1.05-2.32)** | 1.45 (0.95-2.20) |
| Q4 (7.50-15.83) | **6.91 (4.41-10.80)\*** | **4.35 (2.60-7.29)\*** | **3.35 (2.16-5.20)\*** | **2.51 (1.55-4.05)\*** | 1.42 (0.91-2.23) | 1.31 (0.81-2.12) |
| P for trend | <0.001 | <0.001 | <0.001 | <0.001 | 0.090 | 0.218 |
| **Body shape index (ABSI)**□ |  |  |  |  |  |  |
| Q1 (0.0643-0.0803) | 1.00 | 1.00 | 1.00 | 1.00 | 1.00 | 1.00 |
| Q2 (0.0804-0.0840) | 1.55 (0.99-2.42) | 1.59 (0.96-2.63) | 1.31 (0.89-1.95) | 1.33 (0.87-2.04) | 0.93 (0.63-1.37) | 0.91 (0.60-1.38) |
| Q3 (0.0841-0.0874) | **3.55 (2.27-5.56)\*** | **4.03 (2.37-6.86)\*** | **1.81 (1.19-2.76)** | **1.99 (1.23-3.21)** | 1.46 (0.96-2.20) | 1.40 (0.88-2.22) |
| Q4 (0.0875-0.1034) | **3.17 (2.06-4.88)\*** | **3.17 (1.86-5.38)\*** | 1.30 (0.86-1.97) | 1.33 (0.87-2.04) | 0.95 (0.63-1.44) | 0.87 (0.54-1.39) |
| P for trend | <0.001 | <0.001 | 0.115 | 0.240 | 0.846 | 0.691 |

OR: Odds ratio; CI: Confidence interval; W: Women; M: Men.
<sup>†</sup>Adjusted for gender, age, educational level, alcohol consumption, smoking, skin phenotype, vitamin D supplementation, season of blood collection and cognitive function and frailty status.
Waist circumference was further adjusted for height. Bold text indicates a statistically significant difference with a p-value less than 0.05.
□ Missing data in 10 individuals.
\* $p \leq 0.001$

In the second quartile of serum 25(OH)D levels, there was also a significant positive association with the third BRI quartile (OR: 2.14; 95% CI: 1.37-3.34) and fourth BRI quartile (OR: 2.51; 95% CI: 1.55-4.05), *P* for trend <0.001. Similarly, the third ABSI quartile was also positively associated with second quartile of serum 25(OH)D levels (OR: 1.99; 95% CI: 1.23-3.21).

Additionally, when an interaction effect between frailty status and adiposity indices was tested statistical differences were not found.

## Discussion

In this cross-sectional study an inverse association between frailty and obesity with serum 25(OH)D concentrations, independently of sex, age, educational level, alcohol consumption, smoking, skin phenotype, vitamin D supplementation, season of blood collection and cognitive function, was found. These results were consistent with findings of several meta-analyses which evaluate the association between each of these conditions with vitamin D deficiency (18,20,21). The interaction between frailty and adiposity indices concerning serum 25(OH)D was further explored but no significant results were found, meaning that frailty and obesity are independently associated with lower serum 25(OH)D levels, and the effect of frailty (or obesity) on serum 25(OH)D levels is the same at all levels of obesity (or frailty).

When we compared vitamin D levels between frailty and obesity groups we found decreasing 25(OH)D concentrations across them. Thus, individuals that were not frail or obese presented higher unadjusted median 25(OH)D serum concentrations than the other study participants. Interestingly, when data were stratified by sex the pattern was very similar in women, however results were not as evident among men. These observations were supported after by the results of logistic regression, which revealed an association between these conditions and lower serum 25(OH)D levels.

All adiposity indicators evaluated were inversely associated with 25(OH)D serum concentrations. Regarding BMI, an inverse association between 25(OH)D levels and obesity, but not for pre-obesity was found. Moreover, being at the fourth quartile of BRI was associated with a four-fold increased risk of presenting 25(OH)D levels in the first quartile, and it was more strongly associated than the other studied adiposity indicators. It was also observed that the odds of being in the first quartile of 25(OH)D increased significantly across BRI quartiles.

Physiological changes that occur with ageing predispose older adults to lower levels of serum 25(OH)D and frailty status. In addition, lower vitamin D concentrations may also have a negative impact on frailty status through multiple pathways. It has been previously demonstrated that vitamin D was linked with physical function, muscle strength and physical activity (25,43,44). Accordingly, results from several clinical trials carried out in older adults showed that vitamin D supplementation had a beneficial effect in muscle strength and function (15). Evidence suggests the presence of vitamin D receptors (VDR) in the muscle which mediate multiple effects (45). Furthermore, several mechanisms have been suggested to explain the association between muscle function and vitamin D deficiency. In more depth, vitamin D may play an important role in muscle, mediated by several signaling pathways derived from genomic and non-genomic actions of VDR. These mechanisms include regulation of calcium homeostasis, cell proliferation and differentiation, fibers size and protection against insulin resistance, fatty degeneration of the muscle and arachidonic acid mobilization (46). Nevertheless, this receptor was recently found to be undetectable in skeletal muscle, which brings this issue to the fore (47). On the other hand, frailty may contribute to lower 25(OH)D levels, since frail older adults may spend fewer hours engaged in outdoor activities and, consequently, have a reduced sunlight exposure.

Present study results are also consistent with previous data reporting an inverse relationship between 25(OH)D levels and increased adiposity (20–22,28,29). Besides vitamin D deficiency being frequent in older adults, it is also common in obese people. A possible explanation is that obese individuals usually have less skin exposed compared with normal weight individuals (48). Nevertheless, we were unable to evaluate sunlight exposure in the present research. A study which intended to explore the causality and direction of this association using genetic markers, revealed that a higher BMI leads to lower 25(OH)D concentrations (49). In addition, improvement in circulating levels of 25(OH)D was observed in pre-obese and obese after a weight loss intervention (50,51). Since adipose tissue acts as a reservoir for vitamin D, it has been hypothesized that inadequate levels of vitamin D in obese individuals may be predisposed by the sequestration of vitamin D by fat tissue (52). However, it has been recently suggested that this association may be related to a simple volumetric dilution due to higher volume of distribution of 25(OH)D in the adipose tissue (53). Therefore, it is expected that individuals with higher levels of adiposity may be predisposed to inadequate serum 25(OH)D concentrations. Supporting the volumetric dilution hypothesis, a higher dose was required to produce the desired increment in serum 25(OH)D concentrations among obese individuals (54). This supports the Endocrine Society guidelines, which state that the therapy should be adjusted in the presence of obesity (1). Also, evidence suggests that adipocytes express VDR (55), 25-hydroxylase (56) and 1α-hydroxylase enzymes (56,57) which are involved in vitamin D metabolism. Interestingly these enzymes seem to have a decreased expression in obesity (56).

Body mass index and WC are traditionally chosen as anthropometric indicators of general and abdominal adiposity, respectively. Nevertheless, in the present study, the other adiposity indices evaluated (BRI and ABSI), were positively associated with lower vitamin D levels, showing that these may also be used as alternative obesity indicators to identify older adults at risk of low 25(OH)D levels. Despite the lack of positive correlation between ABSI and BMI, our study also demonstrated the link between these indices and lower vitamin D levels, which reinforces their utility.

The present study has some limitations. Firstly, this was a cross-sectional study, therefore the possibility of reverse causation should not be excluded. Secondly, although we have adjusted for multiple covariates, the possible occurrence of residual confounding cannot be ruled out. Thirdly, serum 25(OH)D concentrations were measured using electrochemiluminescence immunoassay, when liquid chromatography-tandem mass spectrometry is considered the golden standard, which can introduce variability in the results (58). And, lastly, participants′ sun exposure levels were not assessed.

In contrast, some strengths can also be pointed out. To our knowledge, this is the first study to explore the association of BRI and ABSI with serum 25(OH)D levels and to elucidate the impact of both obesity and frailty status on 25(OH)D serum levels. Moreover, for all the studied sample, vitamin D was dosed with the same method, the same equipment and in the same laboratory. The very low serum 25(OH)D levels in our sample, with only 30% of the sample presenting adequate 25(OH)D serum concentrations, allowed to study this association.

In summary, present results show that besides BMI and WC, other measures of adiposity such as BRI and ABSI are inversely associated with 25(OH)D serum concentrations. As discussed above, several studies reported conflicting results, however present results reinforce the positive relationship between vitamin D deficiency and both frailty and obesity. Plus, they emphasize the need to target obese and frail elderly people and monitoring their serum vitamin D levels with special care. However, longitudinal studies are necessary to fully elucidate these associations.

---

**Supplementary Table 1. Comparison of the included and excluded individuals^†^.**

**Supplementary Table 2. Characteristics of the 1447 study participants by obesity and frailty status.**

**Supplementary Table 3. Correlation between body mass index, waist circumference, body roundness index and body shape index.**

